# Oropharyngeal mucosal transmission of Zika virus in rhesus macaques

**DOI:** 10.1101/107128

**Authors:** Christina M. Newman, Dawn M. Dudley, Matthew T. Aliota, Andrea M. Weiler, Gabrielle L. Barry, Mariel S. Mohns, Meghan E. Breitbach, Laurel M. Stewart, Connor R. Buechler, Michael E. Graham, Jennifer Post, Nancy Schultz-Darken, Eric Peterson, Wendy Newton, Emma L. Mohr, Saverio Capuano, David H. O’Connor, Thomas C. Friedrich

## Abstract

Zika virus (ZIKV) is present in urine, saliva, tears, and breast milk, but the transmission risk associated with these body fluids is currently unknown. We evaluated the risk of ZIKV transmission through mucosal contact in rhesus macaques. Application of high-dose ZIKV directly to the tonsils of 3 rhesus macaques resulted in detectable plasma viremia in all animals by 2 days post-exposure; virus replication kinetics were similar to those observed in animals infected subcutaneously. Three additional macaques inoculated subcutaneously with ZIKV served as saliva donors to assess the transmission risk from contact with oral secretions from an infected individual. Seven naive animals repeatedly exposed to donor saliva via the conjunctivae, tonsils, or nostrils did not become infected. Our results suggest that there is a risk of ZIKV transmission via the mucosal route, but that the risk posed by oral secretions from individuals with a typical course of ZIKV infection is low.

## Introduction

Zika virus (ZIKV) is a mosquito-borne flavivirus that is associated with Guillain-Barré syndrome in adults and a range of birth defects, most notably microcephaly, in congenitally infected infants^1–5^. ZIKV has become a subject of global concern as it has rapidly expanded its geographic range in the past 2 years. One of the many surprising aspects of the current ZIKV pandemic is the confirmation of one previous report of sexually transmitted ZIKV infection^6–9^. Because mosquito-borne flavivirus infection has not previously been associated with human-to-human transmission, understanding transmission risks is critical for designing effective prevention and control strategies.

In the Americas, ZIKV is transmitted among humans by *Aedes* species mosquitoes, primarily *Aedes aegypti*^10–13^. The explosive epidemic in tropical areas where important vector mosquito species are common has made it challenging to differentiate between vector-borne and sexual ZIKV transmission, so the risk of human-to-human transmission has been difficult to assess. In the continental United States, which has primarily travel-associated cases, detection of non-vector transmission is much more straightforward, and 41 cases of sexually transmitted ZIKV infection have been reported as of January 27, 2017(https://www.cdc.gov/zika/geo/united-states.html). These cases have included male-to-female, male-to-male, and female-to-male transmission^6,14–16^.

ZIKV RNA has been detected in blood, semen, and vaginal secretions, consistent with observations of sexual transmission. Viral RNA and/or infectious virus has also been reported in urine, saliva, tears, and breast milk, suggesting that these body fluids may also pose a transmission risk^17–19^. Indeed, virus was transmitted to a caregiver of an individual with high ZIKV viremia who eventually succumbed to infection^20^. Here we used rhesus macaques to evaluate the risk of ZIKV transmission through mucosal membrane contact with saliva (specifically, palatine tonsils, nasal mucosae, and conjunctivae). We applied high-dose ZIKV stock directly to the tonsils of one group of animals to assess what we term the “theoretical” risk of mucosal transmission using a dose that is >20-fold higher than that typically found in saliva. In another experiment, we applied saliva from ZIKV-infected macaques to determine whether ZIKV might also be transmitted by more casual contact.

## Results

### ZIKV application to tonsils results in systemic infection

To evaluate the risk of ZIKV transmission via the oropharyngeal mucosa, we applied 8 × 10^5^ PFU Zika virus/H.sapiens-tc/FRA/2013/FrenchPolynesia-3328 (ZIKV-FP;^21^) directly to the palatine tonsils of three Indian-origin rhesus macaques using a pipet (Fig. 1). One macaque infected by this route (664817) had detectable plasma viremia by 1 day post-infection (dpi) and all 3 had detectable plasma viremia by 2 days post-infection (dpi) (Fig. 2a). Peak plasma viremia was observed at 6 dpi in all 3 animals and ranged from 2.4 × 10^5^ to 1.1 × 10^7^ vRNA copies/mL. ZIKV plasma viremia was undetectable in all animals by 14 dpi. Plasma viremia of animals infected after tonsillar inoculation (Fig. 2, orange traces) was compared to plasma viremia of 3 animals infected subcutaneously (Fig. 2, blue traces). Overall, plasma viremia in macaques infected via the oral mucosa was similar in magnitude and duration to that of macaques inoculated subcutaneously with the same stock of ZIKV-FP. Two notable exceptions were that viremia only became detectable two days after infection in 2 of the 3 macaques infected via the tonsils, and the peak plasma viral load in these animals occurred 1-3 days later than in animals infected subcutaneously (Fig. 2a).

**Figure 1.**
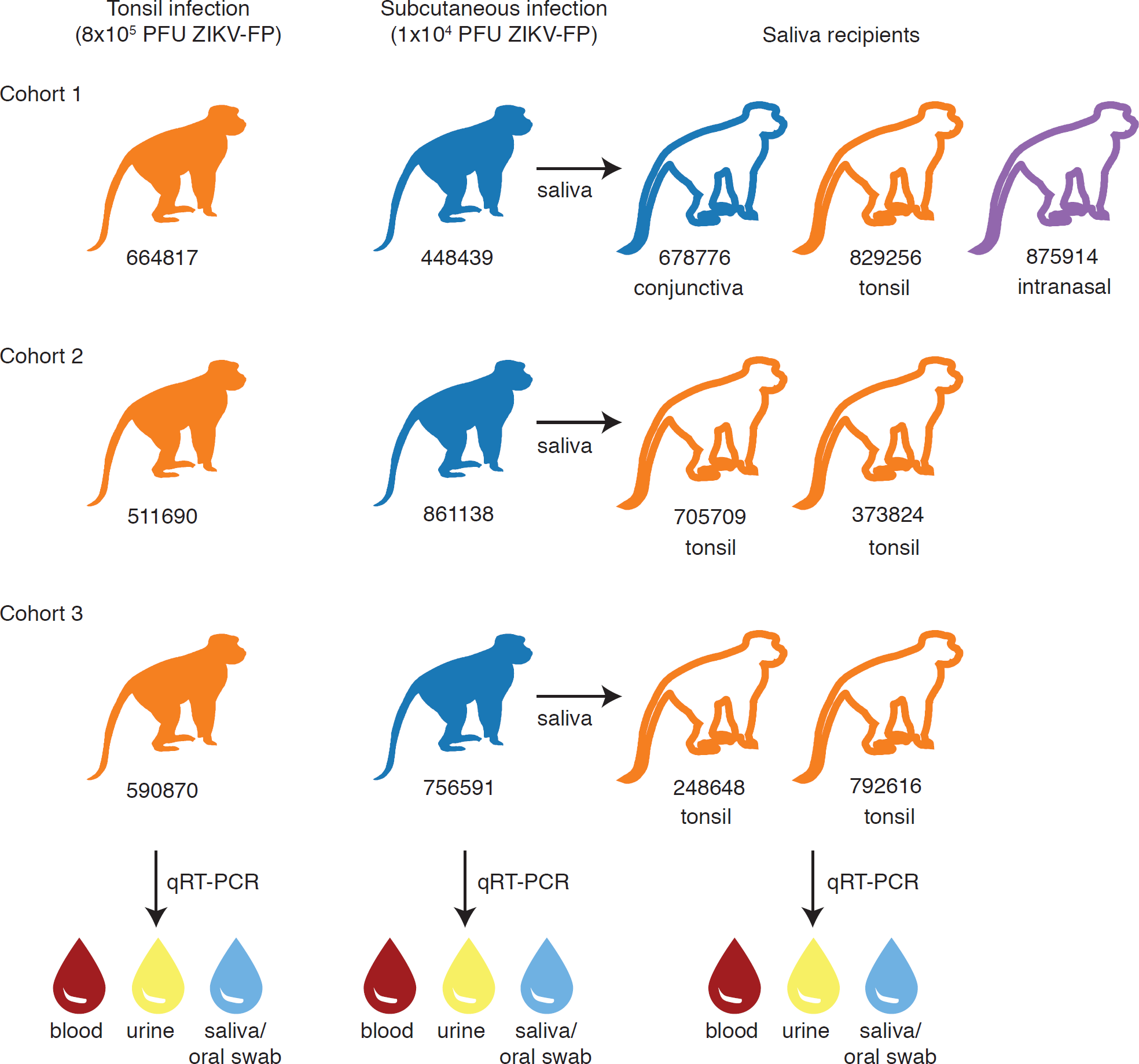
Study design. Three cohorts of animals were inoculated with Zika virus either by application of stock virus to the tonsils (orange filled symbols) or subcutaneously (blue filled symbols). Saliva from animals infected subcutaneously was used to challenge naive recipient animals (open symbols) either to the palatine tonsils, conjunctivae or nasal passages. Blood plasma, urine and oral swabs (and/or saliva) were tested for Zika virus RNA by qRT-PCR in all animals.

**Figure 2.**
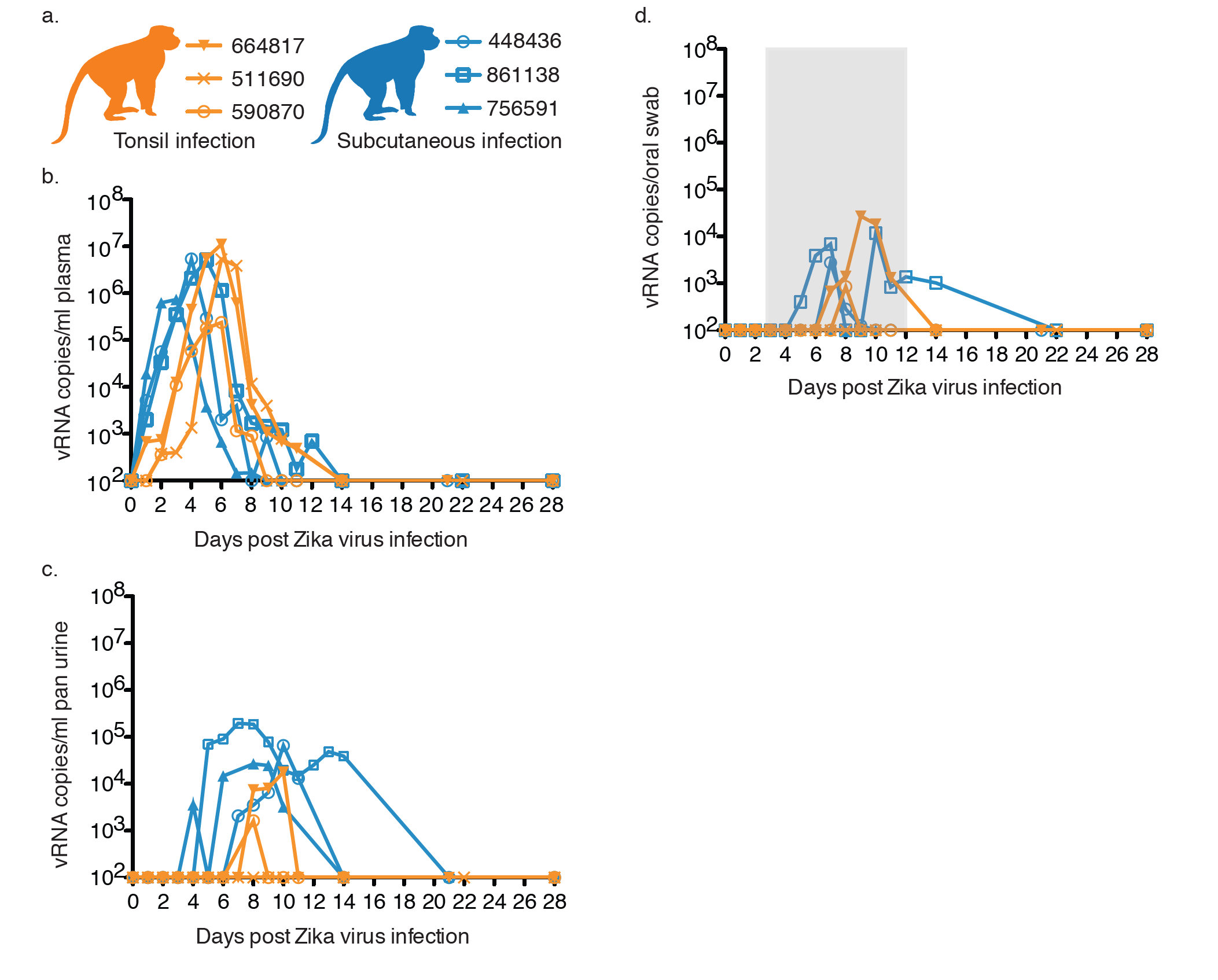
Longitudinal detection of Zika vRNA in plasma, urine and oral swabs in subcutaneously infected animals (blue) or animals inoculated directly to the tonsils (orange). a. Legend for graphs represented in parts b-d. Orange lines represent animals infected via the tonsil and blue lines represent animals infected via subcutaneous injection. b. Zika vRNA copies per mL of peripheral blood plasma. c. Zika vRNA copies per mL of passively collected urine from pans. d. Zika vRNA copies per oral swab. The gray box indicates the time frame in which saliva or an oral swab sample from saliva donor animals was used to challenge recipient animals. The y-axis starts at the limit of quantification of the qRT-PCR assay (100 vRNA copies/mL).

Zika virus RNA was also detectable in other bodily fluids following tonsil exposure. ZIKV RNA was detected in the urine of animal 664817 at 8 dpi (1.6 × 10^3^ vRNA copies/mL) and 590870 at 8-10 dpi (range = 7.4 × 10^3^ - 1.8 × 10^4^ vRNA copies/mL), but not in 511690 (Fig. 2b). We collected saliva when possible, but because the volume of saliva that could be collected was often insufficient to accurately assess viral load, we also collected oral secretions using absorbent swabs that were eluted in a standardized volume of viral transport media (VTM). ZIKV RNA detection in oral swab samples was variable from most animals and undetectable from animal 511690 (Fig. 2c). We were only able to collect saliva directly on two occasions from animal 664817, and both samples were negative for ZIKV RNA (data not shown). ZIKV RNA was detected in saliva from animal 590870 on days 6-9 post infection and from animal 511690 on days 8 and 10 post infection (Fig. 3). Overall, when saliva samples were available, ZIKV RNA was detected more consistently and at more time points than in oral swabs (Fig. 3). By 14 dpi, ZIKV RNA was undetectable in the tested body fluids of all animals infected by tonsil inoculation.

**Figure 3.**
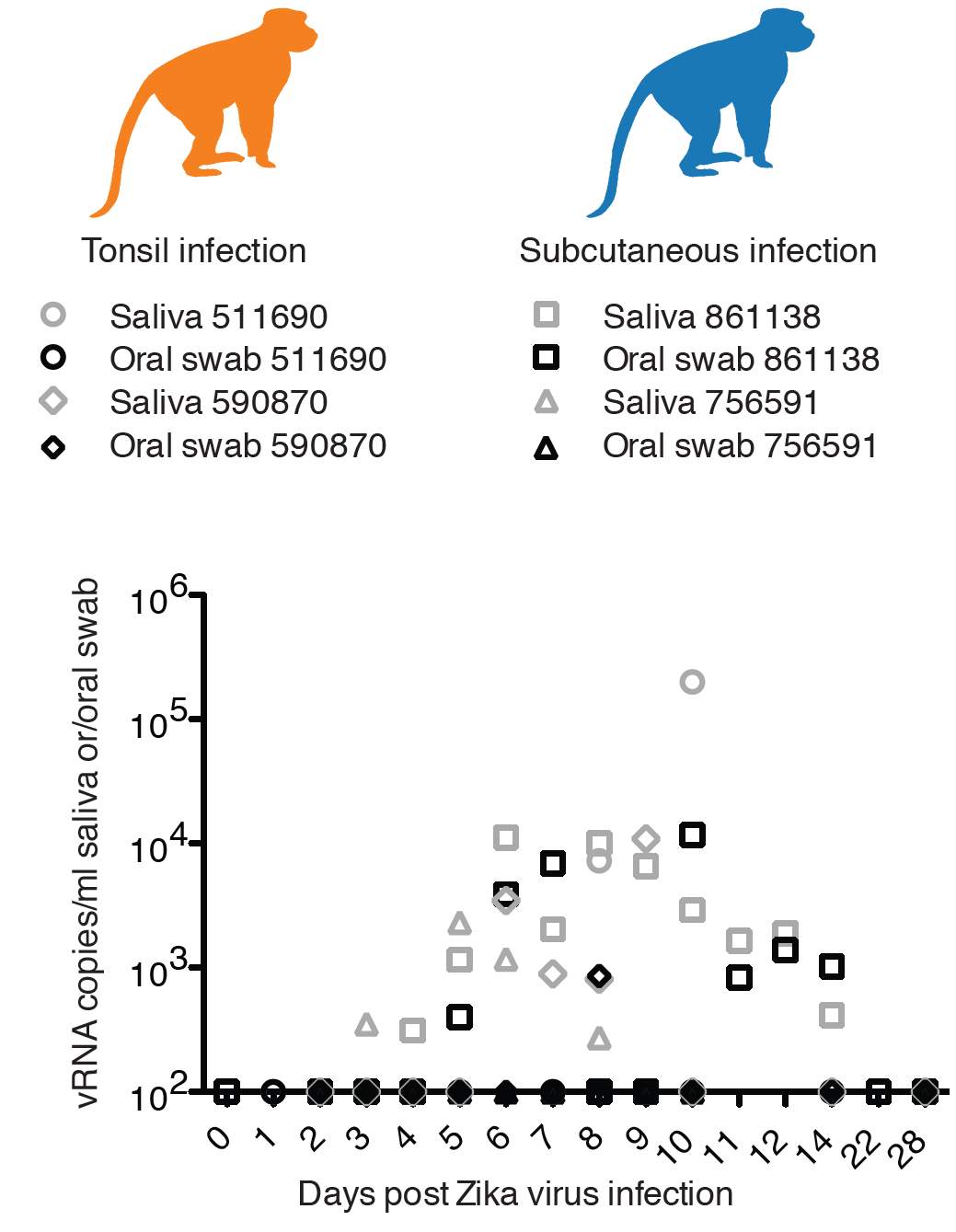
Longitudinal Zika virus load detected in saliva and oral swabs. Two of three animals infected either via the tonsil (under orange monkey) or subcutaneously (under blue monkey) had sufficient saliva collected for viral load testing. Only time points in these four animals at which both saliva (gray) and oral swabs (black) were tested simultaneously are shown. All points not distinguishable above the limit of quantification (100 vRNA copies/mL) are present on the x-axis.

Sera from macaques that were infected by application of ZIKV to the tonsils neutralized ZIKV-FP across a range of serum dilutions. Indeed, neutralization curves prepared using sera from all 3 animals revealed a similar profile as compared to sera from animals infected subcutaneously (see Supplementary Figure 1). All animals developed neutralizing antibodies (nAb) with a 90% plaque reduction neutralization test (PRNT_90_) titer of 1:160 (664817 and 511690) or 1:80 (590870) by 28 dpi. -Together, these results show that application of high doses of ZIKV to the oral mucosae can result in systemic infection and induce humoral immune responses in a manner similar to subcutaneous infection.

### Inoculation with oral secretions from ZIKV-infected donors does not result in systemic infection

The highest concentration of ZIKV RNA we observed in the oral secretions of infected macaques in previous studies was 2.9 × 10^4^ vRNA copies/mL, while the dose of ZIKV stock applied to the tonsils of the animals described above contained 8 × 10^5^ PFU (approximately 8 × 10^8^ vRNA copies/mL). To examine whether saliva from ZIKV-infected macaques was infectious and represented an actual transmission risk to uninfected macaques through oropharyngeal mucosa exposure, we first subcutaneously inoculated 3 rhesus macaques with 1 × 10^4^ PFU of ZIKV-FP, using the same dose, stock, and strain we have reported previously^21,22^. These animals served as saliva donors to 7 naive recipients—saliva or oral swabs were collected from donors daily from 3-10 dpi (cohorts 1 and 3) or 3-12 dpi (cohort 2) and applied to the tonsils, nostrils, or conjunctivae of 1 or more recipients (See Fig. 1 for details). This 3-10 or 3-12 dpi timeframe encompassed the period of time in which ZIKV RNA was detected in non-blood body fluids of animals infected subcutaneously in our previous studies. All saliva donors had detectable ZIKV plasma viremia by 1 dpi, peaking 3-5 dpi, (range = 7.35 × 10^5^ - 5.42 × 10^6^ vRNA copies/mL) and resolving by 14 dpi (Fig. 2a). ZIKV RNA was detected in the passively collected urine of all 3 donors. Peak urine viral loads were detected 7-10 dpi (range = 2.68 x 10^4^ - 1.95 x 10^5^ vRNA copies/mL) (Fig. 2b).

We monitored ZIKV RNA in saliva over time through collection of oral swabs and, whenever possible, saliva, from donor animals, to measure vRNA in the inocula used to challenge the recipients (Fig. 2c and Fig. 3). We detected ZIKV RNA in the oral swab eluate of two saliva donors (448436 and 861138); levels peaked at 2.77 × 10^3^ vRNA copies/mL at 7 dpi and 1.18 × 10^4^ vRNA copies/mL at 10 dpi respectively. The last saliva donor (756591) did not have detectable ZIKV RNA in oral swab samples. We were unable to consistently collect saliva from 448436, but ZIKV RNA in the saliva of the 861138 was detectable 4-14 dpi and peaked at 1.12 × 10^4^ vRNA copies/mL (Fig. 3). 756591 had detectable ZIKV RNA in saliva that peaked at 2.32 × 10^3^ vRNA copies/mL at 5 dpi. Considered together, all donor animals had at least 2 time points with detectable vRNA in either the saliva or oral swab samples during the time when saliva was used to challenge the recipient animals. At 28 dpi, serum collected from all 3 donor animals yielded PRNT_90_ values of 1:160, similar to the recipients of tonsillar challenges (see Supplementary Figure 1 for neutralization curves). Importantly, ZIKV RNA was undetectable in all 7 recipient animals in all body fluids tested throughout the study. Furthermore, ZIKV-specific neutralizing antibodies were undetectable in recipient animals ≥ 21 days after the final challenge with saliva from infected donors.

### Infectivity of saliva from subcutaneously infected donors to IFNAR -/- mice

Because norecipients of saliva from infected donors had detectable systemic ZIKV infections, we tested the infectivity of donor saliva in *IFNAR-/-* mice. These mice are highly susceptible to ZIKV infection^23,24^, even at challenge doses as low as 1 × 10^2^ fluorescent focus units^23^ (approximately 1 × 10^2^ PFU (personal communication Michael Diamond)). We inoculated 5− to 6-week-old *IFNAR-/-* mice with ZIKV RNA-positive macaque saliva from 861138 collected at 10 dpi (1.12 x 10^4^ vRNA copies/mL; 50 μl inoculum; ~1 × 10^−0.25^ PFU (estimated from 1:1000 ratio of PFU:vRNA copies/mL of ZIKV stock ^21,22^)) or oral swab eluate from 448436 collected at 7 dpi (2.77 × 10^3^ vRNA copies/mL; 50 μl inoculum; ~1 × 10^−0.86^ PFU (estimated)), and monitored them for 25 days for weight loss and mortality. All control and experimental mice survived with no evidence of morbidity (Supplementary Fig. 2). Furthermore, ZIKV RNA was undetectable in mouse serum at 3 dpi (the approximate peak of viremia in mice infected previously^23,24^) and the animals did not seroconvert by 25 dpi, as assessed by PRNT (data not shown). These data suggest that saliva from ZIKV-infected macaques did not contain enough replication-competent virus to initiate infection in these sensitive hosts. The same saliva and oral swab samples were concurrently inoculated onto Vero cells to attempt to quantify the infectious dose the mice might have received; however, both the saliva and oral swab sample used for the mouse challenge experiments gave no plaques on these cells. Our plaque assay and mouse inoculation doses contained on the order of 100-500 vRNA copies, while our previous studies suggest that the ratio of vRNA copies to plaque-forming units is approximately 1000:1, further supporting the conclusion that very little infectious virus was present in these samples^21,22^. Together these results suggest that macaques with ZIKV virus loads in the usual range after subcutaneous infection shed very small quantities of infectious ZIKV in saliva.

## Discussion

Humans and animals infected with ZIKV are known to shed virus (or vRNA) in multiple body fluids, including blood, urine, saliva, and genital secretions^21,25–27^. Sexual transmission of ZIKV between humans has been documented in several cases, but the risk of transmission from more casual contact has been difficult to evaluate. Human-to-human transmission via non-sexual contact has been reported in a single case to date, though this case involved a source patient with extremely high viremia; in this case contact with tears or sweat from the source patient was hypothesized to be the mode of transmission^20^. Here we used a nonhuman primate model to investigate whether exposure to ZIKV via mucous membranes, particularly the oropharyngeal mucosa, represents an infection risk.

Application of a high dose (8 × 10^5^ PFU) of Asian-lineage ZIKV to the palatine tonsils resulted in systemic infection in 3 of 3 rhesus macaques, suggesting that productive infection can indeed be initiated at the oropharyngeal mucosa. The kinetics and magnitude of ZIKV replication in plasma in these three animals were similar to those observed in animals infected subcutaneously with the same stock of ZIKV-FP (Fig. 2a), and all 3 animals developed strong nAb titers against homologous ZIKV-FP by 28 dpi. Detection of ZIKV RNA in oral swabs, saliva, and urine was variable in the animals that received a direct tonsil challenge, findings that were also similar to those from animals infected subcutaneously in this and a previous study^21^.

Limitations associated with sample collection could have influenced our ability to detect vRNA in fluids other than blood. For example, time from urination to sample collection varied because urine was collected passively from pans in the animals’ housing. Similarly, there is wide variation in the degree to which individual animals salivate while under sedation. As a result, it was not always possible to collect saliva from each animal at each time point. Oral swabs provided a more consistent means for collecting oral secretions, but as the relatively small amount of secretions absorbed by the swabs must be eluted in medium, detection of ZIKV RNA or infectivity may not be as sensitive in swab sample as in undiluted saliva. Accordingly, ZIKV RNA was more consistently detected in the saliva versus oral swabs when both sample types were available (Fig. 3). Although ZIKV RNA was detectable in the saliva of ZIKV donor animals 861138 and 756591 and in oral swab samples from 861138 and 448436, transfer of saliva from donors to the mucosae of naive recipients did not result in ZIKV transmission. In addition, donor saliva produced no detectable plaques on Vero cells. Moreover, *IFNAR*-/- mice inoculated with saliva or oral swab samples from infected donor macaques showed no overt signs of disease and did not seroconvert. Our previous studies suggest that the proportion of vRNA copies to infectious particles from both virus stock and sera from infected animals is approximately 1000:1^21,22^. The highest viral load that we detected in a saliva sample in this study was only 1.1 × 10^4^ vRNA copies/mL; given that we were never able to collect a full 1mL of saliva, it seems likely that saliva transferred to recipients (tonsil maximum of 100-200μL, conjunctivae maximum of 50μL, and nasal passage maximum of 100μL) in our study likely contained less than three infectious ZIKV virions.

In addition to sample collection limitations, saliva may represent a natural barrier to virus transmission in the oral cavity. A number of different viruses, such as HIV and influenza A, are detected in the saliva of infected individuals, but mucosal infection in many cases is considered low risk^28–30^. This may be due in part to the antimicrobial constituents present in saliva, which include mucins, lysozyme, peroxidase, defensins, and salivary agglutinin^30^. However, even with these natural barriers to infection, challenge of animals with a higher dose of ZIKV directly to the tonsils resulted in productive infection. A bolus challenge of SIV directly to the oral cavity has also been shown to infect rhesus macaques^31^. The authors of that study suggested that their oral bolus transmission model might approximate HIV transmission through oral-genital contact^31^. The finding that an oral bolus of virus may overcome natural mucosal barriers to infection is especially interesting because ZIKV has been detected in semen for many months following acute infection and at viral loads significantly higher than those detected in blood plasma (8.6 log_10_ copies/mL)^32,33^. ZIKV RNA and infectious virus have also been detected in breast milk, another potential route of oral mucosal exposure^34^. The potential for transmission of HIV through breast milk is well documented in humans and is associated with the milk viral load^35,36^. Although there have been no documented cases of transmission between a nursing mother and her infant, a ZIKV viral load of greater than 8 × 10^6^ vRNA copies/mL has been reported in the breast milk in a woman with acute ZIKV infection^18^.

Taken together, our results suggest that there is a risk of ZIKV transmission via the oral mucosal route, as shown by systemic infection in 3 of 3 macaques after application of high-dose infectious virus to the tonsils. However, the actual risk of transmission via mucosal exposure to saliva may be low—saliva from donor animals with typical plasma viremia harbored little or no infectious virus and could not mediate transmission of ZIKV in our study. However, it must be noted that secretions, including saliva from individuals with unusually high viral load, semen, and breast milk, could pose a transmission risk.

## Materials and methods

### Study Design

This study was designed as a proof of concept study to examine whether ZIKV transmission may occur in the absence of vector-borne or sexual transmission, via saliva, in the rhesus macaque model. Nothing is currently known about the potential saliva transmission of ZIKV in vivo so we selected 3 animals for direct application of ZIKV stock to the palatine tonsils, 3 animals for subcutaneous inoculation with a well characterized dose of ZIKV^21^ to serve as saliva donors, and 7 animals (a minimum of 2 per donor) to serve as recipients of saliva from infected donors. Available animals were allocated to experimental groups based on qualitative assessment of salivation while under ketamine sedation as communicated by staff at the Wisconsin National Primate Research Center. Investigators were not blinded to experimental groups.

### Ethical approval

This study was approved by the University of Wisconsin-Madison Institutional Animal Care and Use Committee (Animal Care and Use Protocol Number G005401 and V5519).

### Nonhuman primates

Six male and seven female Indian-origin rhesus macaques (*Macaca mulatta*) utilized in this study were cared for by the staff at the Wisconsin National Primate Research Center (WNPRC) in accordance with the regulations, guidelines, and recommendations outlined in the Animal Welfare Act, the Guide for the Care and Use of Laboratory Animals, and the Weatherall report. In addition, all macaques utilized in the study were free of Macacine herpesvirus 1, simian retrovirus type D, simian T-lymphotropic virus type 1, simian immunodeficiency virus, and had no history of exposure to any dengue virus serotype. Animals ranged in age from 3 years to 17 years old (mean = 7.8 years). For all procedures, animals were anesthetized with an intramuscular dose of ketamine (10mL/kg). Blood samples were obtained using a vacutainer or needle and syringe from the femoral or saphenous vein.

### Virus

Macaques in this study were inoculated with Asian-lineage Zika virus/H.sapiens-tc/FRA/2013/FrenchPolynesia-01_v1c1 (ZIKV-FP) obtained from Xavier de Lamballerie (European Virus Archive, Marseille, France). This virus was originally isolated from a 51-year-old female in France after travel to French Polynesia in 2013 and passaged a single time on Vero cells (African green monkey kidney cells; CCL-81). The virus stock used in this study was prepared by inoculation onto a confluent monolayer of C6/36 mosquito cells (*Aedes albopictus* larval cells; CRL-1660). Cell lines were obtained from American Type Culture Collection (ATCC), were not further authenticated, and were not specifically tested for mycoplasma. A single, clarified harvest of virus, with a titer of 5.9 × 10^6^ PFU/mL (3.9 × 10^9^ vRNA copies/mL) ZIKV-FP was used as stock for all subcutaneous inoculations.

### Tonsil challenges

The ZIKV-FP stock was thawed, diluted in PBS to 1 × 10^6^ PFU/mL, and maintained on ice until inoculation. Each animal was anesthetized and 8 × 10^5^ PFU ZIKV-FP was applied directly to the palatine tonsils (maximum of 400μL per tonsil) via pipet after visualization with a laryngoscope. Animals were closely monitored by veterinary and animal care staff for adverse reactions and signs of disease. Animals were examined, and blood, urine, oral swabs, and saliva were collected from each animal daily from 1 through up to 12 dpi and then weekly thereafter through 28 dpi.

### Subcutaneous inoculations

The ZIKV-FP stock was thawed, diluted in PBS to 1 × 10^4^ PFU/mL, and loaded into a 3mL syringe maintained on ice until inoculation. For subcutaneous inoculations, each of three Indian-origin rhesus macaques was anesthetized and inoculated subcutaneously over the cranial dorsum with 1mL virus stock containing 1 × 10^4^ PFU. All animals were closely monitored by veterinary and animal care staff for adverse reactions and signs of disease. Animals were examined, and blood, urine, oral swabs, and saliva were collected from each animal daily from 1 through up to 12 dpi and then weekly thereafter through 28 dpi.

### Mucosal membrane challenges

Oral swabs or saliva were collected from subcutaneously inoculated, ZIKV-infected donor macaques for mucosal membrane challenge of uninfected recipient animals beginning at day three post-infection and continuing through day 10 (cohorts 1 and 3) or 12 (cohort 2) post-infection of the donor animals. For recipients challenged with saliva from the first donor, oral swabs were held under the tongue of the anesthetized donor animal for up to 2 minutes to collect saliva. Swabs were then placed in 750μL of viral transport media (tissue culture medium 199 supplemented with 0.5% FBS and 1% antibiotic/antimycotic), swished vigorously for at least 15 seconds, and then swabs were discarded. Viral transport media containing saliva was then applied via pipet to the palatine tonsils (100μL per tonsil) (after visualization via laryngoscope), conjunctivae (50μL per eye), or nasal passages (100μL per nostril) of the corresponding recipient animal. For recipients challenged with saliva from the second and third donors, saliva was collected via pipet from under the tongue of an anesthetized donor animal and applied directly via pipet to the palatine tonsils (200μL per tonsil) of each recipient animal. Mucosal challenges were repeated daily for up to 10 days. Animals were closely monitored by veterinary and animal care staff for adverse reactions and signs of disease. As described previously, animals were examined, and blood, urine, saliva, and oral swabs were collected from each animal daily during challenge and then weekly thereafter through 28 days after final challenge.

### Plaque reduction neutralization test (PRNT)

Macaque serum samples were screened for ZIKV neutralizing antibody utilizing a plaque reduction neutralization test (PRNT) on Vero cells (ATCC #CCL-81). Endpoint titrations of reactive sera, using a 90% cut off (PRNT_90_), were performed as described in^37^ against ZIKV-FP. Neutralization curves were generated using GraphPad Prism software. The resulting data were analyzed by non-linear regression to estimate the dilution of serum required to inhibit 50% of infection.

### Live virus isolation

Mice deficient in the alpha/beta interferon receptor on the C57BL/6 background (*IFNAR-/-*) were bred in the pathogen-free animal facilities of the University of Wisconsin-Madison School of Veterinary Medicine. Four groups (n=12) of 5- to 6-week-old, mixed sex mice were used to test infectivity of ZIKV RNA in macaque saliva. Mice were inoculated in the left, hind foot pad with 50μL of saliva or saliva swab eluate in viral transport media (VTM) from samples that tested positive for ZIKV RNA by qRT-PCR. Mock-infected experimental control mice received saliva or saliva swab eluate in VTM collected from ZIKV-negative control macaques. Following inoculation, mice were monitored twice daily for the duration of the study. Sub-mandibular blood draws were performed and serum was collected to verify viremia via qRT-PCR and nAb titers via PRNT.

### Viral RNA isolation

Plasma was isolated from EDTA-anticoagulated whole blood collected the same day by Ficoll density centrifugation at 1860 rcf for 30 minutes. Plasma was removed to a clean 15mL conical tube and centrifuged at 670 rcf for an additional 8 minutes to remove residual cells. Urine was opportunistically collected from a pan beneath each animal’s cage and centrifuged at 500 rcf for 5 minutes to remove cells and debris. Saliva was collected via pipet from under the tongue or using sterile oral swabs run under the tongue while animals were anesthetized. Saliva collected via pipet was used to directly inoculate recipient animals in cohorts 2 and 3. Additional saliva, when available, was removed from the collection pipet, centrifuged at 500 rcf for 5 minutes, and added to 650μL viral transport media at a ratio of no more than one part saliva to three parts viral transport media. Swabs were placed in viral transport media for 60-90 minutes, then vortexed vigorously and centrifuged at 500 rcf for 5 minutes to elute saliva. Prior to extraction, oral swab eluates were pelleted by centrifugation at 14000 rpm and 4ºC for an hour. After centrifugation, supernatant was removed, leaving virus in 200μL media. Viral RNA was extracted from 300μL plasma or urine using the Viral Total Nucleic Acid Kit (Promega, Madison, WI) on a Maxwell 16 MDx instrument (Promega, Madison, WI). Viral RNA was extracted from 200μL oral swab-derived samples using the QIAamp MinElute Virus Spin Kit (Qiagen, Germantown, MD) with all optional washes. RNA was then quantified using quantitative RT-PCR. Viral load data from plasma and urine are expressed as vRNA copies/mL. Viral load data from oral swabs are expressed as vRNA copies/mL eluate. Viral load data from saliva in viral transport media are expressed as vRNA copies/mL total volume.

### Quantitative reverse transcription PCR (qRT-PCR)

For ZIKV-FP, vRNA from plasma, urine, saliva, and oral swabs was quantified by qRT-PCR using primers with a slight modification to those described by Lanciotti et al. to accommodate African lineage ZIKV sequences^38^. The modified primer sequences are: forward 5’-CGYTGCCCAACACAAGG-3’, reverse 5’-CACYAAYGTTCTTTTGCABACAT-3’, and probe 5’-6fam-AGCCTACCTTGAYAAGCARTCAGACACYCAA-BHQ1-3’. The RT-PCR was performed using the SuperScript III Platinum One-Step Quantitative RT-PCR system (Invitrogen, Carlsbad, CA) on a LightCycler 480 instrument (Roche Diagnostics, Indianapolis, IN). The primers and probe were used at final concentrations of 600 nm and 100 nm respectively, along with 150 ng random primers (Promega, Madison, WI). Cycling conditions were as follows: 37°C for 15 min, 50°C for 30 min and 95°C for 2 min, followed by 50 cycles of 95°C for 15 sec and 60°C for 1 min. Viral RNA concentration was determined by interpolation onto an internal standard curve composed of seven 10-fold serial dilutions of a synthetic ZIKV RNA fragment based on ZIKV-FP.

### Data availability

Primary data that support the findings of this study are available at the Zika Open-Research Portal (https://zika.labkey.com). The authors declare that all other data supporting the findings of this study are available within the article and its supplementary information files, or from the corresponding author upon request.

## Acknowledgements

We thank the Veterinary, Animal Care and Scientific Protocol Implementation staff at the Wisconsin National Primate Research Center (WNPRC) for their contribution to this study. We thank Emma Walker for assistance with the mouse studies. We acknowledge Jens Kuhn and Jiro Wada for preparing silhouettes of macaques used in figures. We thank the DHHS/PHS/NIH R01AI116382 to D. H. O. for funding. We also thank the P51OD011106 awarded to the WNPRC, Madison-Wisconsin. This research was conducted in part at a facility constructed with support from Research Facilities Improvement Program grants RR15459-01 and RR020141-01. The publication’s contents are solely the responsibility of the authors and do not necessarily represent the official views of NCRR or NIH.

### Author Contributions

T.C.F., C.M.N., D.M.D., M.T.A., and D.H.O. designed the experiments. C.M.N., D.M.D., M.T.A., and T.C.F analyzed data and drafted the manuscript. M.T.A. provided and prepared viral stocks, performed plaque assays, and performed the mouse experiments. A. M. W., G. L. B. and T.C.F. developed and performed viral load assays. M.S.M., 3M.E.B., L.M.S., C.R.B, C.M.N., E.L.M and D.M.D. coordinated and processed macaque samples for distribution. M.E.G. maintained the Zika Open Portal site where data was stored and shared. J.P., N.S-D., E.P., S.C., and W.N., coordinated and performed macaque infections and sampling.

### Competing financial interests

The authors declare no competing financial interests.

### Materials and Correspondence

Thomas C. Friedrich

Department of Pathobiological Sciences, University of Wisconsin-Madison, Madison, WI 53706

Virology Services, Wisconsin National Primate Research Center, University of Wisconsin-Madison, Madison, WI 53715

---

**Supplementary Figure 1. Neutralization by ZIKV immune sera from tonsil inoculated and subcutaneously inoculated macaques.** Immune sera from macaques inoculated subcutaneously (blue) or via the tonsils (orange) were tested for their capacity to neutralize ZIKV-FP. ZIKV was mixed with serial 2-fold dilutions of serum for 1 hour at 37°C prior to being added to Vero cells. Infection was measured by plaque reduction neutralization test (PRNT) and is expressed relative to the infectivity of ZIKV-FP in the absence of serum. The concentration of sera indicated on the x-axis is expressed as Log_10_ (dilution factor of serum). The dilution of sera at half-maximal neutralization of infection (EC_50_) was estimated by non-linear regression analysis and were 2.803 (861138), 3.081 (448436), 2.479 (756591), 2.638 (664817), 2.991 (511690), and 2.746 (590870). Neutralization curves for each animal at 28 dpi are shown.

**Supplementary Figure 2. Injection of ZIKV+ oral swab eluate or ZIKV+ saliva does not cause mortality or morbidity in *IFNAR*-/- mice.** *IFNAR*-/- mice (n=3 per group) were inoculated in the left hind footpad with ZIKV+ or ZIKV-oral swab eluate (black) or ZIKV+ or ZIKV-saliva (gray). Mice were monitored until 25 dpi; all survived without signs of morbidity. Changes in weight were calculated daily for ZIKV+ and ZIKV-oral swab eluate and ZIKV+ or ZIKV-saliva inoculated mice. Error bars represent the standard deviation of the mean.

